# Using herbarium samples for NGS methods – a methodological comparison

**DOI:** 10.1101/2021.08.26.457828

**Authors:** Pia Marinček, Natascha D. Wagner, Salvatore Tomasello

**Affiliations:** Department of Systematics, Biodiversity and Evolution of Plants (with Herbarium), University of Goettingen, Untere Karspüle 2, 37073 Göttingen, Germany

## Abstract

Herbaria harbor a tremendous amount of plant specimens that are rarely used for plant systematic studies. The main reason is the difficulty to extract a decent quantity of good quality DNA from the preserved plant material. While the extraction of ancient DNA in animals is well established, studies including old plant material are still underrepresented. In our study we compared the standard Qiagen DNeasy Plant Mini Kit and a specific PTB-DTT protocol on two different plant genera (*Xanthium* L. and *Salix* L.). The included herbarium material covered about two centuries of plant collections. A selected subset of samples was used for a standard library preparation as well as a target enrichment approach. The results revealed that PTB-PTT resulted in higher quantity and quality regarding DNA yield. For relatively recent herbarium specimens, and despite the lower overall yield of DNA, the Qiagen Kit resulted in better sequencing results regarding the number of filtered and mapped reads. We were able to successfully sequence a sample from 1820 and conclude that it is possible to include old herbarium specimens in NGS approaches. This opens a treasure box for phylogenomic research.

## Introduction

Molecular biodiversity research as well as phylogenomic studies rely on a good, comprehensive sampling. However, very frequently the required material is either not available, e.g., in case of extinct species, or not accessible, e.g., if species occur in very remote areas. To overcome the problems of insufficient sampling, herbarium specimens could be used as a source of information (1,2). Herbaria harbor a massive amount of specimens that were collected over several centuries and can thus be treated as treasure troves for biodiversity research (3–7). It is estimated that around 70,000 new species are already housed in herbaria, “waiting to be described” (6). However, although herbarium vouchers are a valuable source of information, using them for molecular studies remained challenging (2,8).

The DNA of herbarium samples is usually highly degraded and fragmented and extracting DNA from old tissues remains difficult. Mainly because of both, generally limited success of DNA extraction and the challenges associated with PCR-amplification of highly degraded DNA, researchers avoid to include historical specimens (2). In Sanger sequencing times, amplification and sequencing required long, intact DNA fragments, and therewith incorporating historical samples, especially from plants, was almost impossible. In contrast, more recent developments in sequencing techniques enabled researchers to include fragmented DNA (=short fragments) in their approaches (7,9). Nevertheless, a certain level of DNA quality and quantity is necessary to include historical material in studies using NGS methods.

For most phylogenomic studies, the DNA is usually extracted from fresh or silica dried plant material by using a commercial DNA extraction kit. Historical samples require more advanced methods with special regard to shorter fragment length and putative contamination (10). However, extracting DNA from plant cells is per se more complicated than from animal cells, especially for historical samples. Weiß et al. (11) found out that plant DNA in herbaria showed a six times higher fragmentation rate than animal DNA preserved in bones. The high number of secondary compounds, including polyphenolics and polysaccharides that can covalently bind to DNA or coprecipitate with it, are known to inhibit PCR even in non-degraded DNA samples. This complicates the usage of DNA from plant herbarium tissues (7,12). Additionally, the quality and quantity of DNA found in herbarium specimens depends on conditions during collection and storage, and is, in general, lower than for freshly collected plant material followed by immediate drying in silica gel or freezing (1,13,14).

The first studies on ancient or archival DNA (aDNA) from plants were published in the early 90s of the last century and dealt with plant remains in archaeological sites (e.g., (15,16) among others). Studies dealing with DNA extraction from old herbarium samples have used one of the following approaches: i) Early studies on herbarium material aiming at sequencing single markers, e.g., ITS, simply used standard CTAB protocols for extraction (17) or a modified version of it (12,18–21); ii) commercial kits were used with few adaptations, e.g., increasing incubation times (18,19,22,23); or, iii) more specific protocols for aDNA extraction were applied (e.g., optimized to obtain short sequences and to increase the proportion of endogenous DNA in the extracts; (10,12,13,24). Since then, more and more studies included historical plant material in phylogenomic studies (7,23,25,26). However, specific protocols for aDNA are generally more expensive, time consuming and require specific facilities and hygiene rules, not always available in systematic botany laboratories. Moreover, extraction protocols were often optimized for a certain taxonomic group or model organism (27).

Although a few recent studies focused on comparing the efficiency of CTAB-based extraction protocols with commercial kits (19), or comparing CTAB extractions with protocols specific for aDNA (10), no studies yet have investigated the circumstances in which aDNA methods (e.g., the PTB-DTT protocol described in (28)) should be preferred to commercial kits when extracting DNA from old and damaged herbarium material. For systematists (systematic botanists) extraction kits represent still the easiest and most convenient solution for DNA extraction.

In the study presented here, we want to test in which circumstances it is recommendable to invest more time and resources to extract DNA from herbarium specimens using a specific aDNA protocol (PTB-DTT) instead of a standard kit. We measure DNA yield and quality of the PTB-DTT approach and the standard Qiagen DNeasy Plant Mini Kit on herbarium material of different age and condition. Additionally, we want to test, whether the resulting DNA can be used for standard NGS library preparation (i.e., double stranded library preparation for Illumina sequencing), and target enrichment approaches using commercially available kits. To incorporate the taxonomic effect on extraction performance, we performed the analyses using specimens from different taxonomic groups of two phylogenetically very distant plant genera.

## Material and Methods

### Plant material

To test the different extraction methods, we used herbarium material of two distinct plant genera, i.e., *Salix* L. and *Xanthium* L. For genus *Salix* we included three species with each four samples: *S. caprea*, a diploid tree or big shrub that is frequently distributed in central Europe, *S. myrsinifolia*, a widely distributed hexaploid tree and *S. breviserrata*, an alpine diploid dwarf shrub. The herbarium samples were collected in the herbarium Göttingen (GOET) and covered about two centuries. The oldest herbarium sheet was from 1820, the youngest from 2015.

For *Xanthium*, we included samples from the two sections of the genus, i.e., section *Xanthium* (plants with unarmed stems) and section *Acanthoxanthium* DC. (plants with spiny stems). Specimens were from the herbarium Göttingen (GOET), from the herbarium of the Botanic Garden and Botanical Museum Berlin-Dahlem (B), and from herbarium of the Bavarian Natural History Collections (M), with the oldest being collected in 1821 and the youngest in 1984. We used in total 25 *Xanthium* accessions. For details of all samples used in this study, see Table 1.

**Table 1.**
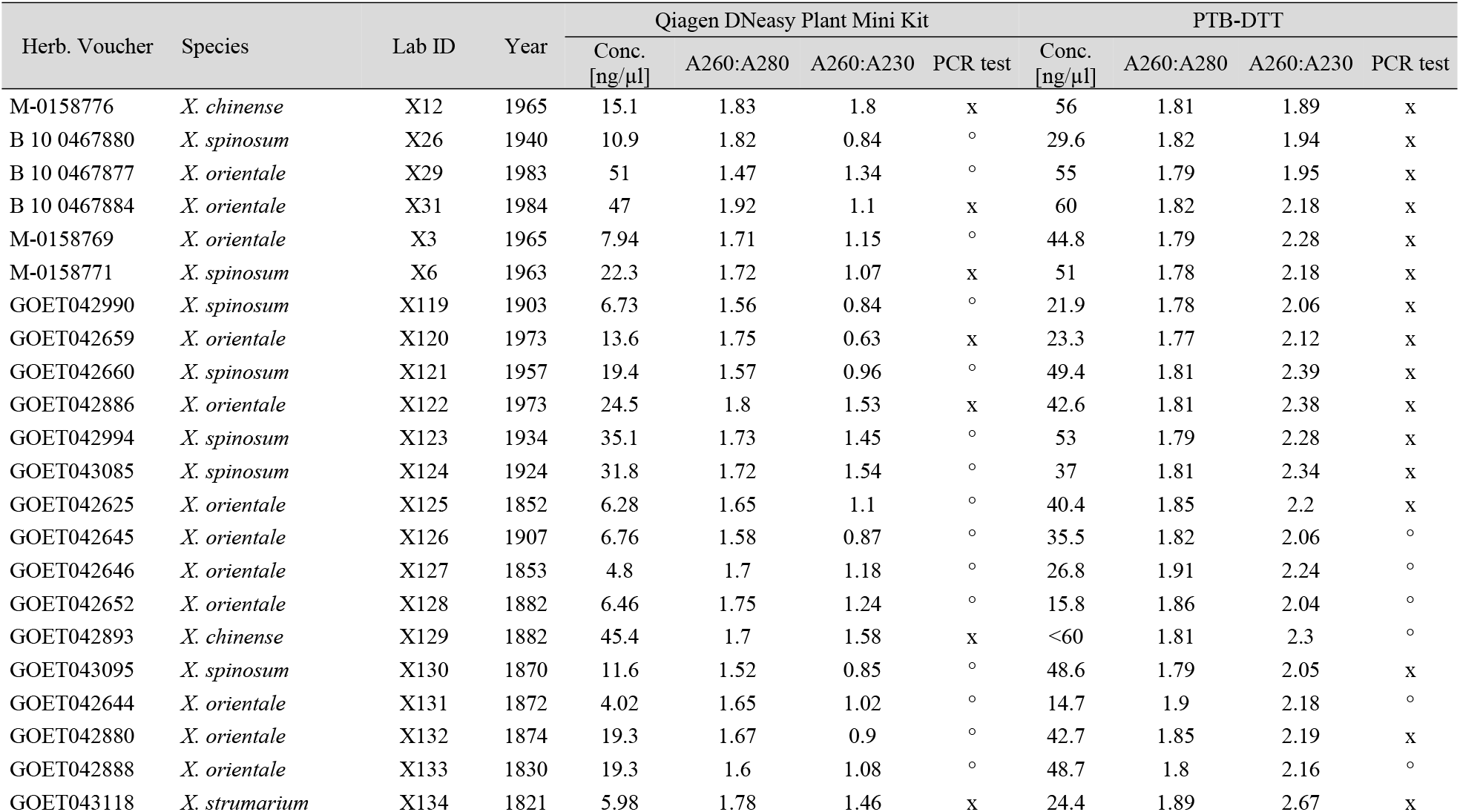
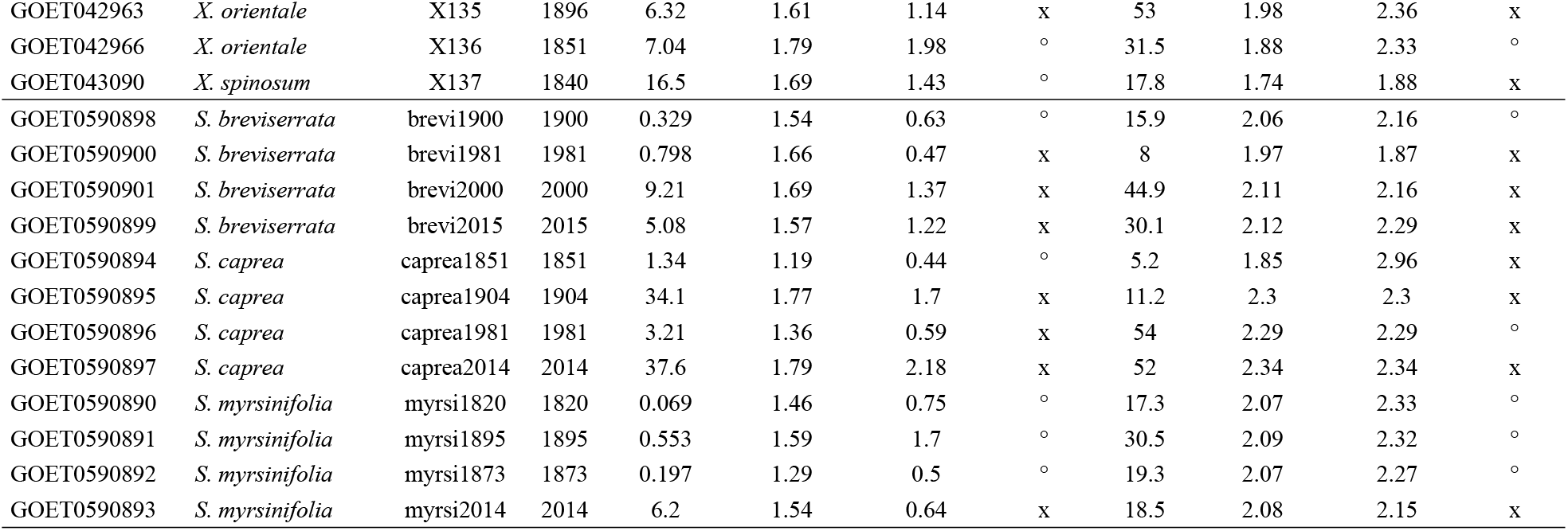
Information on the herbarium specimens of *Xanthium* and *Salix* used in this study. It includes year of collection, concentrations, and absorbance ratios’ values for the Qiagen DNeasy Plant Mini Kit and the PTB-DTT extractions. Successful PCR amplifications are indicated by the symbol *x*, PCR failures by °. Species assignment in *Xanthium* follows (30).

### DNA extraction

For each sample about 10 mg of leaf material was removed from the herbarium sheet and transferred to an Eppendorf tube. The material was pulverized using a TissueLyser II (Qiagen, Venlo, Netherlands). PTB-DTT extractions were done as described in Dabney et al. (29), and following the modifications applied by Gutaker et al. (28). Additionally, the DNA of all samples was extracted using the Qiagen DNeasy Plant Mini Kit according to the manufacturer’s instructions (Qiagen, Venlo, Netherlands) and with the following modifications: i) lysis incubation as well as the incubation on ice after adding the P3 buffer were prolonged to 30 minutes (instead of ten and five minutes, respectively); ii) during DNA elution, 50 μl of AE buffer (instead of 100 μl) were added to the column and incubated for 30 minutes (instead of five) before centrifugation. The elution step was then repeated resulting in 100 μl suspension of DNA in elution buffer.

All extractions were performed under hygienical precautions typical for working with aDNA. Surfaces and consumables were sterilized with DNA AWAY (ThermoFisher Scientific, Waltham, US) and pipets were UV-treated before and after each extraction using a nUVaClean™ UV Pipette Carousel (MTC Bio, Metuchen, US). Extractions were carried out under a laminar flow hood wearing mask and full-body laboratory suits.

### DNA yield and quality measurements

Since the same amount (10 mg) of herbarium material was employed in each extraction, we used concentrations as measure of DNA yield. Concentrations were measured on a Qubit 3 fluorometer (Thermo Fisher Scientific, Waltham, US), using the Qubit dsDNA HS assay kit (Thermo Fisher Scientific, Waltham, US). To measure the A260:A280 and A260:A230 absorbance ratios, we used a Nanodrop 2000 (Thermo Fisher Scientific, Waltham, US).

Additionally, we ran electrophoresis gels to visually check success of the extractions and approximate fragment lengths. We mixed 5 μl of extract with 1μl of Roti^®^-Load DNAstain 3 (Carl Roth, Karlsruhe, Germany), and loaded it in a 2% agarose gel. Two ladders were used to fully cover the fragment length spectrum (long fragments for the recent herbarium specimens, very short ones for the old specimens): the DNA-Ladder 20 bp extended (bands from 20 to 1000 bp; Biozym, Hessisch Oldendorf, Germany) and the 1 kbp DNA-Ladder (bands from 500 to 1000 bp; Carl Roth, Karlsruhe, Germany). Electrophoreses were run for 40 minutes at 100 volts.

### Statistics

To test for correlation between age of the herbarium specimen and DNA yield, we performed Pearson’s correlation tests (31), treating samples from the two genera as well as the two extraction methods separately. ANCOVA was performed to test the effect of the extraction method (Qiagen kit vs. PTB-DTT), and of the taxonomy on the DNA yield (DNA concentration), and quality (A260:A280 and A260:A230 absorbance ratios), treating the voucher age as covariate. We tested ANCOVA assumptions for normality and homoscedasticity with the Levene’s test (32). All statistical analyses as well as scattered- and box-plots were done in R (R Core Team 2018).

### PCR test

As an additional quality check, the extracted DNA was used to amplify the plant plastid locus *trn*L-*trn*F with the primers *e* and *f* (33). 1μl of each sample was mixed with 12.5 μl of Roti^®^-Pol TaqS Master mix (Carl Roth, Karlsruhe, Germany), 1μl of forward and 1μl reverse primer in the concentration of 5pmol/μl each. Finally, 9 μl of sterile, distillated water were added to each sample solution to achieve a final volume of 25 μl. We used a touchdown protocol for amplification with the following settings: denaturation at 94 °C for 2 min, followed by 10 cycles each starting with 20 seconds at 94 °C, 20 seconds at 63°C with a drop of 1°C for each cycle, and 30 seconds at 72°C. Then, 25 cycles followed, starting with 20 seconds at 94°C, followed by 20 seconds at 52°C and 30 seconds at 72°C. The final extension was at 72°C for 5 minutes. To check the amplification success, 1 μl of the PCR product was mixed with 4 μl of ddH_2_O and 1 μl of Roti^®^-Load DNAstain 3 (Carl Roth, Karlsruhe, Germany), and loaded together with the DNA-Ladder 20 bp extended (Biozym, Hessisch Oldendorf, Germany) onto a 2% agarose gel. Electrophoreses were run for 40 minutes at 100 volts.

### Library preparations and sequencing

To analyze to which extent extracts were usable for NGS sequencing and, to estimate the amount of endogenous DNA (i.e., percent or reads mapping to a reference), we sequenced a subset of 12 samples (six *Salix* and six *Xanthium* from both extraction methods) with the Illumina technology (Illumina Inc., San Diego, USA). Libraries were prepared using either the “NEBNext Ultra II DNA Library Prep Kit for Illumina^®^” (for old herbarium specimens) or the “NEBNext Ultra II FS DNA Library Prep Kit for Illumina^®^” (for more recent specimens; New England BioLabs, Ipswich, USA). In both cases, we followed the manufacturer’s instructions, with the only modification that the purification following the adapter ligation was done using 1.5 volumes of HighPrep™ beads (MagBio Genomics, Gaithersburg, US) instead of 0.8 volumes, to minimize the loss of ultra-short fragments. Samples were PCR-amplified for 14 cycles and samples-specific dual indices (“NEBNext Multiplex Oligos for Illumina^®^”, E7600; New England BioLabs, Ipswich, USA) were attached to the fragments.

The *Xanthium* samples were applied to a hybrid capture reaction using the commercially available myBaits COS Compositae 1Kv1 kit (Arbor Biosciences Ann Arbor, Michigan, USA). This was done for two reasons: i) we wanted to investigate if libraries were suitable for a hybrid capture reaction. Standard kits have 120 bp long baits and might not efficiently hybridize the ultra-short fragments of very old herbarium specimens. And ii), since no *Xanthium* genome is available, we could use the target regions of the bait kits as a ‘pseudoreference’ for reads mapping, and therefor estimate the hybrid capture success and the proportion of endogenous DNA. Six indexed samples were pooled in equal quantities, dehydrated in a Concentrator Plus (Eppendorf, Hamburg, Germany), and diluted in 7 μL of ddH2O. The pool was enriched using the baits kit and following the manufacturer’s protocol. Hybridization took place for 20 h at 65 °C. Enriched products were PCR-amplified for 14 cycles using the 2X KAPA HiFi HotStart Mix (KAPA Biosystems, Wilmington, USA) and the P7 and P5 adapters as primers. Concentrations were measured on a Qubit 3 fluorometer (Thermo Fisher Scientific, Waltham, US) and fragment length distribution was checked with the QIAxcel (Qiagen, Venlo, Netherlands). *Salix* libraries presented adapter-dimers peaks at around 125 bp and were therefore treated with the BluePippin (Sage Science, Beverly, USA) to select fragments between 140 and 600 bp, using a 2% cartridge and internal standard. Finally, the samples (six *Salix* libraries and the *Xanthium* hybrid capture pool) were pooled equimolarly and paired-end sequenced on an Illumina MiSeq System (Illumina Inc., San Diego, USA) at the NIG Core Unite (University of Gottingen, Göttingen, Germany), using a 2×150 bp (300 cycles) v2 kit.

### Reads quality check, mapping and plastome reconstruction

The resulting reads were quality checked using FastQC (available at: http://www.bioinformatics.bbsrc.ac.uk/projects/fastqc). Sequence adapters were removed and reads were quality-trimmed using Trimmomatic v. 0.32 (34) with default settings. To analyze the percentage of target and off-target reads, the reads of the six *Salix* samples were mapped to the published *Salix purpurea* reference genome (female clone 94006; *Salix purpurea* v5.1, DOE-JGI, http://phytozome.jgi.doe.gov/). The reads of the six *Xanthium* samples were mapped to a reference consisting of the concatenation of the target exon sequences each separated by stretches of 800 Ns. Mapping was performed using the mem algorithm of BWA/0.7.12 (35) with default settings. The quality filtered reads were also used to reconstruct the plastome for each sample. Therefore, the reads were subjected to a reference-based assembly using Geneious vR11 2020.2.4 (http://www.geneious.com; (36)) as described in (37). As references, we used the available plastomes in Genbank, NCBI, for each species, i.e., *S. breviserrata* [MW435421], *S. caprea* [MW435424], *S. myrsinifolia* [MW435439] and, *X. sibiricum* [MH473582], respectively.

## Results

### DNA yield

In total, the DNA of 37 samples was extracted using the PTB-DTT method as well as the standard Qiagen DNeasy Plant Mini Kit. The results of the gel electrophoreses for all extracts are shown in S1 Fig. The observed DNA concentrations were significantly higher in the PTB-DTT extractions (mean = 34.87 ng/μl) than for the extractions using the Qiagen kit (mean = 14.7ng/μl) when considering the complete dataset (Paired Student’s t-test, *p* = 2.552×10^-8^; Fig 1A). Results are slightly correlated with the age of the herbarium specimen (Pearson’s *r* = 0.34 and *r* = 0.30 for the PTB-DTT and the Qiagen kit, respectively; Fig 1B). Taxon effect (*Salix* versus *Xanthium*) is also significant (*p* = 0.0096), indicating that concentrations of *Xanthium* DNA extracts (mean = 28.57) were significantly higher than in *Salix* (mean = 16.9).

**Fig 1.**
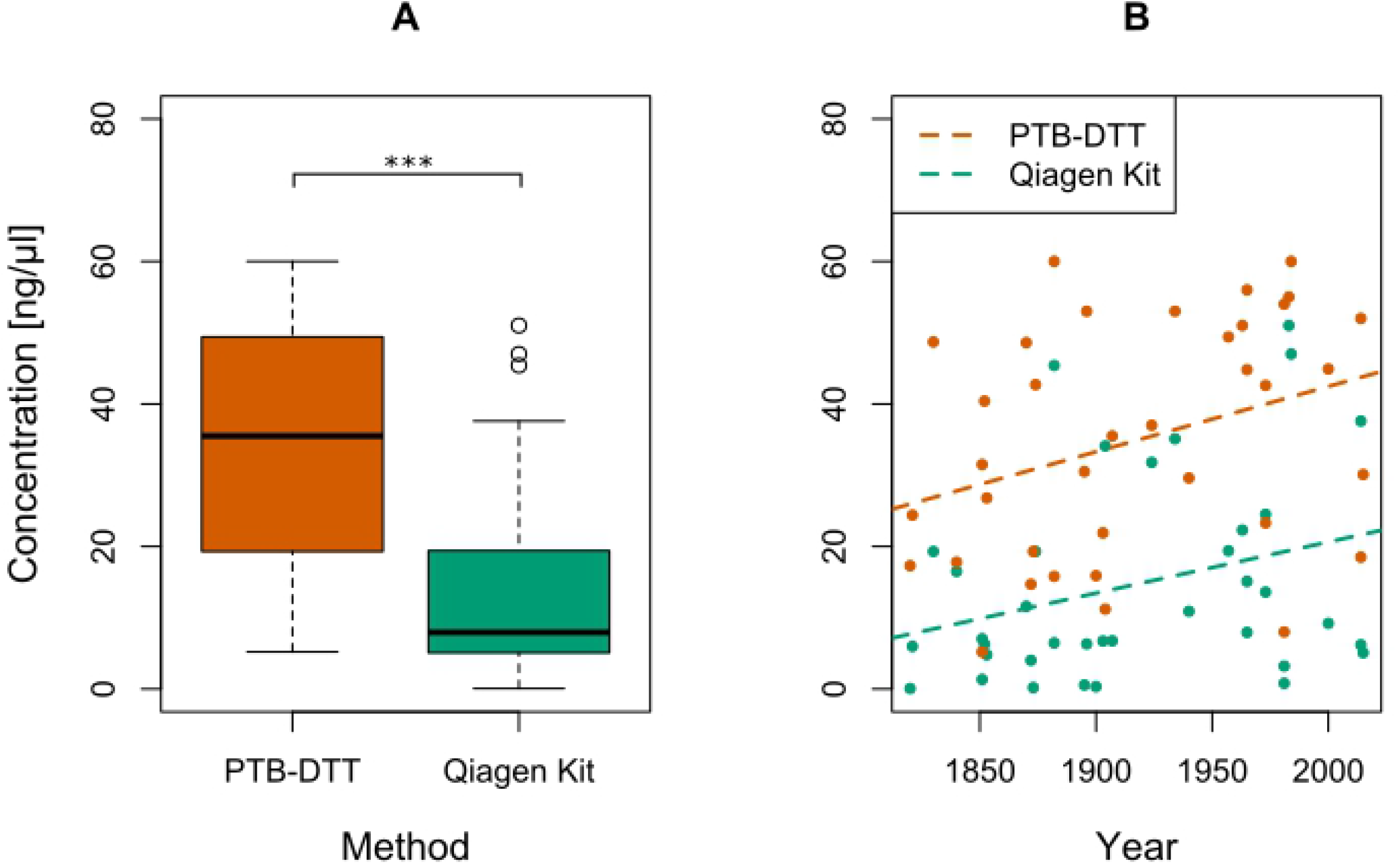
Comparison of the DNA concentration (in ng/μl) of all extracts for both extraction methods. (A) Boxplots of the DNA concentration of all samples extracted with the PTB-DTT and Qiagen Plant Mini Kit extraction protocols. Asterisks represent statistical significance: (*) *p* < 0.05, (**) *p* < 0.01, (***) *p* < 0.001. (B) Scatterplot of the DNA concentrations of all extracted samples against the age of the respective herbarium sheets (year of origin). Lines represent a general linear model for DNA concentration against the year of the herbarium sheet for PTB-DTT and Qiagen kit protocols separately.

When treating the two genera separately, results were similar to those presented above. PTB-DTT extractions performed better than the Qiagen kit (*p* = 4.203×10^-9^ and *p* = 0.007 in *Xanthium* and *Salix*, respectively; see S2 Fig). The taxonomic effect (i.e., differences within different species of *Salix* or sections of *Xanthium*) was neither significant in *Salix (p* = 0.184) nor in *Xanthium* (*p* = 0.909). As for the complete dataset, concentrations were slightly negatively correlated with the age of the specimens, both in *Xanthium* (*r* = 0.43 and *r* = 0.47 in the PTB-DTT and the Qiagen kit, respectively; S3 FigA) and in *Salix* (*r* = 0.56 and *r* = 0.31; S3 FigB).

### DNA quality

A high-quality DNA shows a A260:A280 ratio of 1.8 and a A260:A230 ratio above 2. Our results revealed that the DNA quality was overall higher for the PTB-DTT extractions. The A260:A280 ratios were significantly higher (*p* = 9.5×10^11^) in the PTB-DTT extracts (means = 1.92) compared to the ones obtained with the Qiagen kit (mean = 1.64) (Fig 2A). The results of the A260:A230 ratios could not be statistically compared, because the groups show a significant heterogeneity of variances (Levene test, *p*-value = 0.00012) (Fig 2B).

**Fig 2.**
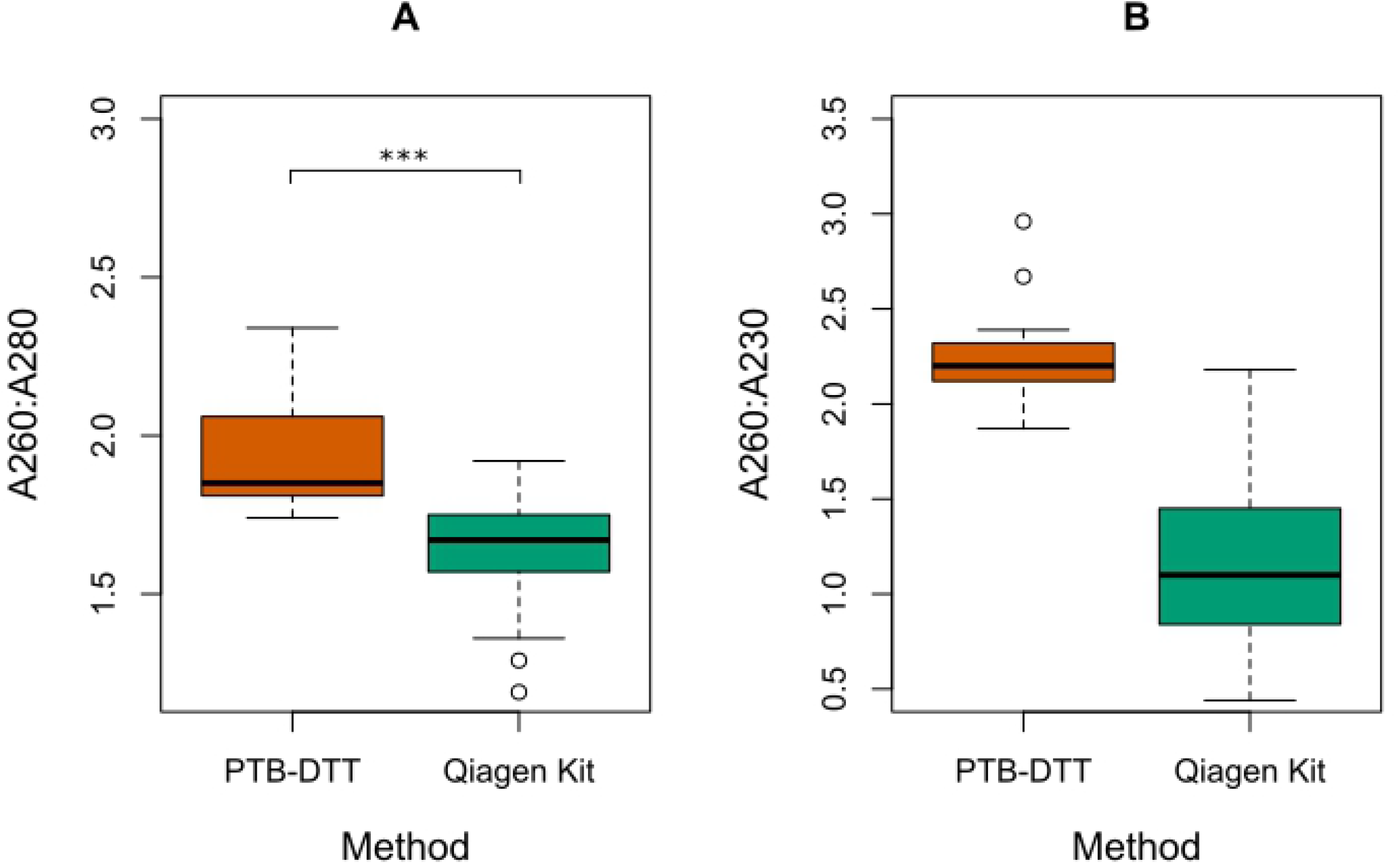
Comparison of DNA quality of all extracts for both tested extraction methods. (A) Comparison of the A260:A280 ratios, measured for all samples for the PTB-DTT as well as the Qiagen DNeasy PlantMini Kit extraction protocols. (B) Comparison of the A260:230 ratios measured for all samples for PTB-DTT and Qiagen kit extraction protocols separately. Asterisks represent statistical significance: (*) *p* < 0.05, (**) *p* < 0.01, (***) *p* < 0.001.

### PCR

The success of the amplification of the plastid *trn*L-*trn*F spacer was assessed by a visible band on the agarose gel. For 25 out of 37 samples of the PTB-DTT extracts the amplification was successful, while 15 out of 37 samples extracted with the Qiagen kit showed amplification success. Regarding the two genera, a total of 26 *Xanthium* samples (out of 50 amplifications) and 14 *Salix* samples (out of 24) were successfully amplified for both extraction methods (see Table 1 for details).

### NGS sequencing results

The sequencing produced 31,899,780 reads in total. On average, we obtained 2,658,315 reads per sample, ranging from 4,254,576 (*X. orientale*, X133 PTB) to 979,024 reads (*X. spinosum*, X137 PTB). The number of filtered low quality reads after trimming differed between the two genera. In *Salix*, the percentage of reads excluded after quality trimming was 13.5% (21.45% - 9.13%) and in *Xanthium* 1.44% (1.85% - 1.06%). The number of duplicate reads was 20.53% (34.5% - 14.98%) in *Xanthium* and 0.82% (1.8% - 0.32%) in *Salix*. The number of reads after quality and duplicate filtering was in average 2,632,716 in *Salix* and 1,709,997 in *Xanthium*. The average percent of mapped reads was 85.08% in *Salix* (89.05% - 78.61%), and 62.6% in *Xanthium* (69.4% - 55.91%) (Table 2).

**Table 2.** Results from the sequencing of the 12 samples selected for the library preparation.

| Sample ID | species | Total # of reads | # quality trimmed reads | % quality trimmed reads | # reads without duplicates | % duplicates | # paired reads without duplicates | genome/targeted regions |  | plastome |  |
| --- | --- | --- | --- | --- | --- | --- | --- | --- | --- | --- | --- |
|  |  |  |  |  |  |  |  | # mapped reads | % of mapped reads | # mapped [paired] reads | % of mapped reads |
| X127 PTB | *X. orientale* | 1,664,846 | 1,634,471 | 1.83 | 1,389,725 | 14.98 | 810,379 | 898,313 | 64.64 | 18,760 | 2.31 |
| X133 PTB | *X. orientale* | 4,254,576 | 4,209,480 | 1.06 | 2,757,444 | 34.5 | 2,087,569 | 1,853,679 | 67.22 | 1,905 | 0.09 |
| X135 PTB | *X. orientale* | 3,035,086 | 3,000,746 | 1.14 | 2,535,678 | 15.5 | 1,488,551 | 1,759,861 | 69.4 | 8,382 | 0.56 |
| X119 PTB | *X. spinosum* | 1,539,976 | 1,511,586 | 1.85 | 1,261,404 | 16.56 | 745,772 | 743,944 | 58.98 | 23,343 | 3.13 |
| X137 PTB | *X. spinosum* | 979,024 | 964,606 | 1.48 | 752,872 | 21.96 | 477,935 | 420,956 | 55.91 | 2,145 | 0.45 |
| X137 QIA | *X. spinosum* | 1,970,678 | 1,945,724 | 1.27 | 1,562,864 | 19.68 | 963,618 | 928,844 | 59.43 | 28,281 | 2.93 |
| 2000 PTB | *S. breviserrata* | 4,207,912 | 3,558,700 | 15.43 | 3,544,188 | 0.41 | 1,507,542 | 2,826,025 | 79.74 | 175,488 | 11.64 |
| 2000 QIA | *S. breviserrata* | 3,059,196 | 2,779,898 | 9.13 | 2,771,084 | 0.32 | 1,302,529 | 2,496,801 | 90.1 | 72,785 | 5.59 |
| 1981 PTB | *S. caprea* | 2,808,198 | 2,402,821 | 14.44 | 2,359,571 | 1.8 | 1,102,771 | 2,015,842 | 85.43 | 89,037 | 8.07 |
| 2014 PTB | *S. caprea* | 3,084,266 | 2,756,045 | 10.65 | 2,743,545 | 0.46 | 1,273,101 | 2,425,974 | 88.42 | 119,865 | 9.42 |
| 2014QIA | *S. caprea* | 2,271,790 | 2,044,809 | 10 | 2,029,125 | 0.77 | 982,885 | 1,827,017 | 90.04 | 88,303 | 8.98 |
| 1820 PTB | *S. myrsinifolia* | 3,024,232 | 2,375,746 | 21.45 | 2,348,788 | 1.14 | 1,138,641 | 1,938,528 | 82.53 | 133,866 | 11.76 |

The plastome assembly was able to recover 100% of the plastome for the three *Salix* species. In detail, 5.59-11.76% of filtered reads mapped to the respective reference plastome. The mean coverage varied between 38 and 104 reads. For both *Xanthium* species, 0.09-3.13% of filtered reads mapped to the reference and between 31% and 83% of the plastome could be recovered. The mean coverage varied between 1 to 210 reads. For more details, see Table 2.

## Discussion

### Effect of specimens’ age on yield and quality

In the presented study, we extracted archival DNA of 37 herbarium specimens, with an age homogeneously spanning 200 years. As in Zeng et al. (38), we found a negative correlation between age of the specimens and DNA yield. Older samples had in general lower yield, especially when using the commercial extraction kit. Our results contrast those of other studies (19,39), where no correlation was found between age of the specimens and DNA yield. The reason of this discrepancy might be explained by sampling peculiarities. In (19), the herbarium specimens employed in their study were not older than 60 years. In Bakker et al. (39), both fresh and herbarium samples were used, most of the latter being not older than 60 years. However, this does not necessarily mean that DNA yield in a very old samples is always lower than in recent herbarium specimens. The extent to which DNA of an old herbarium voucher is degraded depends on other factors for which information is usually scarce (e.g., specimen preparation and conservation conditions). One would expect that plants collected and desiccated in cool and dry environments yield higher quantities of less degraded DNA than plants collected under wet-tropical conditions. Although no studies could fully investigate these aspects yet, Bakker et al. (39) found that, based on reads assembly results, fragmentation effects of the age were more consistent in samples from wet-tropical environments, probably due to longer and more destructive preparation methods (e.g., heat, alcohol).

Moreover, the efficiency of the extraction methods in old specimens may differ considerably in different taxonomic groups (19). In our study, we compared specimens from different taxa of two systematically very distant genera. The negative effect of “aging” was much more pronounced in *Salix* (S3 Fig). When using a standard extraction kit, samples older than 100 years could not produce DNA yield high enough to be employed in standard (double stranded-DNA) library preparation methods (DNA concentrations between 0.069 and 1.34 ng/μl in samples predating 1900; Table 1). On the other hand, in *Xanthium* the Qiagen kit was performing relatively well (in terms of DNA yield) even in samples as old as 200 years.

### Methods performance

Extraction methods specific for old archaeobotanical remains outperform standard extraction methods, in terms of DNA yield and proportion of small endogenous DNA fragments (28). In our study, we confirmed that the PTB-DTT methods produced higher yields compared to widely used extraction kits. In some cases (e.g., for old *Salix* herbarium specimens), using this extraction method was the only mean of gathering enough DNA for library preparation purposes.

Surprisingly enough, the PTB-DTT method outperformed the Qiagen kit also in terms of quality of the DNA extracts (here referred to the absorbance ratios A260:A280 and A260:A230). Our results partially contrast with those of former studies (19), in which the extraction kit (silica-column based) produced purer DNA than the CTAB method. The good performance of the PTB-DTT method could be ascribed here to the fact that the DNA precipitation was also done on a silica-column (as in the Qiagen extraction kit), producing therefore higher quality extracts compared to those obtained with the CTAB protocol in (19). Moreover, in our study the lower quality of the kit extracts could be partially biased by the low absorbance values received by the old herbarium specimens with extremely low DNA concentrations.

The success of the amplification was dependent on the extract quality and concentration. In general, and according to our expectations, the relatively young herbarium specimens performed better than the older ones. A higher number of PTB-DTT extracts produced good amplifications (25 samples) compared to the kit (15 samples). The quality of the extracts (i.e., absence of molecules other than DNA that might eventually inhibit downstream analyses) is particularly important for the success of PCR based techniques (27,40). This was confirmed by the lower success of the PCR amplifications using the kit extractions, especially for the old herbarium specimens. For samples predating 1900, only three and seven PCR reactions were producing bands for the kit and the PTB-DTT extracts, respectively. Regarding the two genera, in *Xanthium* more amplifications were successful than in *Salix*, probably due to the fact that extractions in *Xanthium* generally had better yield and quality than in *Salix*, especially for the old herbarium specimens (see Table 1 for details). Willows are rich in secondary compounds, such as salicylates, tannins or flavonoids (41,42), which unfavorably affects the performance of DNA extractions and downstream molecular analyses.

### Library preparation for Illumina sequencing

We produced libraries for Illumina sequencing for 12 of the 37 samples included in the study, both using PTB-DTT extracts and (when possible) extracts obtained with the Qiagen kit. This was done (i) to test if the extractions produced were quantitatively and qualitatively good enough for library preparation; and (ii) to assess the proportion of endogenous DNA. For the *Salix* samples, libraries were directly sequenced and mapped to a *Salix* reference genome. For *Xanthium*, libraries were enriched using a commercially available baits kit. In this way, we were able to map the obtained reads to the target regions of the baits kit because no reference genome is available for *Xanthium* yet, and to investigate how a commercial kit (non-customized for aDNA) performed on libraries obtained from old herbarium vouchers.

Based on our results we observed a relatively high proportion of low-quality reads in *Salix*, and a high clonality (number of duplicate reads) in *Xanthium*. The former could be attributed to the high number of short and damaged DNA fragments obtained from extractions of old and degraded herbarium vouchers. In *Xanthium* only a small proportion of reads was filtered out due to low quality. Probably, the hybrid-capture reaction helped mitigating this problem by enriching the libraries in those DNA fragments able to bind to the baits (e.g., fragments that were long enough and not too damaged). On the other hand, the number of duplicate reads is relatively high in those samples. Clonality has been reported as a potential problem when target-enrichment techniques are applied to old and damaged DNA (43). This is particularly evident when high numbers of (post capture) PCR cycles are performed on samples with low proportions of endogenous DNA (as potentially old herbarium samples). Increasing the amount of starting DNA (25) or pooling multiple, shorter independent amplifications of a library (43) may help solving this issue. In general, there are a few factors intrinsic of DNA extracted from old and degraded tissues influencing the efficiency of the in-solution hybrid capture reactions (e.g., the low levels of endogenous DNA, the very short DNA fragments;(44)). A few adaptations to the standard protocol may help to partially overcome these problems (e.g., increasing the amount of starting DNA (25) or decreasing hybridization temperature (45)).

In *Salix*, 80%-90% of the reads (after quality filtering) mapped to the reference genome, therefore giving evidence of high proportions of endogenous DNA even in old herbarium specimens. In the oldest sample sequenced, a *S. myrsinifolia* from 1820, about 82% of the reads were mapping to the reference genome. In a similar study (28) only a few samples were able to reach similar percentages of reads mapping to the reference. Our results confirm that standard double-stranded library preparation (as alternative to the more expensive single-stranded library preparation) can produce good and reliable results, especially if the proportion of endogenous DNA in old samples is not extremely low (46). When employing herbarium specimens as old as 200 years, a few adaptations to the protocol may help to optimize the efficiency of dsDNA library preparation (44). It is particularly important to minimize the loss of short endogenous fragments during the purification steps of the library preparation (47). While preparing the libraries, we tried different approaches to minimize the loss of small fragment, especially in the first purification after adapter ligation. We tested (i) the MinElute PCR purification columns (Qiagen, Venlo, Netherlands), capable to retain fragments as short as 70 bp; and (ii) the standard (magnetic beads based) purification with an increased volume of beads (1.5x instead of 0.8x). Given that results from the MinElute and from the modified beads-based purification were comparable, we decided to continue with the latter (more cost-effective) one.

In *Xanthium*, 55%-65% of the reads mapped to the target regions of the baits kit. These proportions are comparable to those obtained using the same kit on fresh (silica-gel dried) samples (data not published). Target enrichment has already been applied successfully on relatively old herbarium specimens (23,25,48). For very old specimens (200 years old and more), methods based on genome skimming and the assembly of multicopy genome regions (e.g., organellar DNA), coupled to single-stranded DNA library preparation, were thought to perform better than target enrichment of single-copy nuclear regions (46). Our results confirm the potential of the latter technique, also when it is applied to herbarium specimens as old as 200 years.

### Plastome assembly

The generated sequencing reads were used to assemble the plastomes of the archival samples. For the six *Salix* samples, between 5.6% and 11.7% of the reads mapped to the respective reference and it was possible to recover the complete plastomes of all samples. The mapping success is in the range of a recent study on *Salix* plastomes based on non-archival, silica dried fresh material. Here, the amount of mapped reads varied between 3.1% and 23.5% (Wagner et al 2021, accepted). For *Xanthium* it was not possible to recover the entire plastome, only 0.1% to 3.1% mapped to the reference and 21-83% of the plastome could be recovered. However, the library preparation differed from the simple skimming approach. The assembly of the plastomes was done based on off-target reads of a target enrichment dataset. In these circumstances, to assemble complete plastomes might be difficult, and focus on the most represented regions could be a valuable alternative (49,50). Nevertheless, our data shown here show the potential to assemble entire plastid genomes from up to 200 years old herbarium samples with standard extraction and sequencing methods. This is in accordance with similar studies (7,9).

## Conclusion

Herbaria harbor huge collections of archival DNA that are still underrepresented in phylogenomic research. Extraction protocols specific for aDNA help to obtain high DNA yields and quality, especially when extracting DNA from old herbarium specimens. However, those methods are usually more expensive and time consuming and require compliance with specific hygiene rules not always feasible in standard systematic botany laboratories. A PTB-DTT extraction takes longer and is more than twice as expensive as a Qiagen DNeasy Plant Mini Kit extraction. Our study showed that it is possible to include herbarium samples from the last two centuries in NGS approaches by using standard commercial DNA extraction, library preparation and target enrichment kits. However, in cases of challenging material (e.g., old samples containing many secondary compounds, as is the case in genus *Salix*) or valuable and rare material (e.g., type material, and/or scarce herbarium sheets) it might be preferable to use specific aDNA extraction protocols.

## Acknowledgements

We thank Dr. Marc Appelhans and Dr. Robert Vogt for support with the herbarium collections in Göttingen and Berlin, respectively.

## Supporting information (Supplements)

**S1 Fig. Electrophoresis gel pictures of *Xanthium* and *Salix* samples extracted with the PTB-DTT protocol and the Qiagen DNeasy PlantMini Kit**.

**S2 Fig. Comparison of DNA concentration (in ng/μl) for both tested extraction methods for both plant genera separately displayed as boxplots.**

(A) DNA concentration of *Xanthium* samples, extracted with the PTB-DTT protocol and Qiagen DNeasy PlantMini Kit. (B) DNA concentration of *Salix* samples extracted with the PTB-DTT protocol and Qiagen kit. Asterisks represent statistically significant differences in means: (*) *p* < 0.05, (**) *p* < 0.01, (***) *p* < 0.001.

**S3 Fig. Comparison of DNA concentration (in ng/μl) for both tested extraction methods for both tested genera separately displayed as scatterplots.**

(A) DNA concentration of *Xanthium* samples extracted with PTB-DTT and Qiagen DNeasy PlantMini Kit protocol against the year of the creation of sample herbarium sheets. (B) DNA concentration of *Xanthium* samples extracted with the PTB-DTT protocol and Qiagen kit against the year of the creation of sample herbarium sheets. Lines represent a general linear model for DNA concentration against the year of the herbarium sheet for the PTB-DTT protocols and Qiagen kit, separately.

